# Associations on the Fly, a new feature aiming to facilitate exploration of the Open Targets Platform evidence

**DOI:** 10.1101/2024.09.10.612089

**Authors:** C Cruz-Castillo, L Fumis, C Mehta, RE Martinez-Osorio, JM Roldan-Romero, H Cornu, P Uniyal, A Solano-Roman, M Carmona, D Ochoa, E McDonagh, A Buniello

## Abstract

**Motivation:** The Open Targets Platform (https://platform.opentargets.org) is a unique, comprehensive, open-source resource supporting systematic identification and prioritisation of targets for drug discovery. The Platform combines, harmonises and integrates data from >20 diverse sources to provide target-disease associations, covering evidence derived from genetic associations, somatic mutations, known drugs, differential expression, animal models, pathways and systems biology. An in-house target identification scoring framework weighs the evidence from each data source and type, contributing to an overall score for each of the 7.8M target-disease associations. However, the previous infrastructure did not allow user-led dynamic adjustments in the contribution of different evidence types for target prioritisation, a limitation frequently raised by our user community. Furthermore, the previous Platform user interface did not support navigation and exploration of the underlying target-disease evidence on the same page, occasionally making the user journey counterintuitive.

**Results:** Here, we describe “Associations on the Fly” (AOTF), a new Platform feature - developed as part of a wider product refactoring project - to enable formulation of more flexible and impactful therapeutic hypotheses through dynamic adjustment of the weight of contributing evidence from each source, altering the prioritisation of targets.

**Availability and implementation:** All Open Targets code is available as open source: [https://github.com/opentargets].

This tool was implemented using React v18 and its code is accessible here: [https://github.com/opentargets/ot-ui-apps].

The tools described in the paper are accessible through the Open Targets Platform web interface [https://platform.opentargets.org/] and GraphQL API (https://platform-docs.opentargets.org/data-access/graphql-api).

Data is available for download here: [https://platform.opentargets.org/downloads] and from the EMBL-EBI FTP: [https://ftp.ebi.ac.uk/pub/databases/opentargets/platform/].

**Supplementary information:** Additional information on this tool can be found on the Platform documentation pages [https://platform-docs.opentargets.org/web-interface, https://platform-docs.opentargets.org/web-interface/associations-on-the-fly, https://platform-docs.opentargets.org/target-prioritisation] and training video [https://youtu.be/2A9bksboAag].

**Contact:** Annalisa Buniello, EMBL-EBI

## Introduction

Open Targets (https://www.opentargets.org/) is a pre-competitive partnership combining expertise from academia and the pharmaceutical industry, aiming to systematically identify and prioritise targets, ultimately leading to safer, more effective therapies. One axis of the consortium’s research programme is to provide open-source data and informatics resources for the global scientific community, with the Open Targets Platform as its flagship tool (https://platform.opentargets.org/)(Ochoa *et al*., 2023).

The Platform harmonises, integrates, and scores publicly available evidence of target-disease associations, mapping to standardised ontologies and identifiers to enable data connectivity and ensure the provenance of the data source. It provides a comprehensive knowledgebase and tooling to identify and prioritise targets, diseases and drugs in the context of drug discovery, as well as annotations of targets, diseases and drugs, including disease causality, target tractability and safety.

The Platform is released four times a year, ensuring regular updates from external data sources and integration of user feedback and new features. As per our latest release (24.03), the Platform includes evidence for 63,226 genes, 25,817 diseases and phenotypes, 17,111 drugs and compounds and 7,802,260 target-disease associations — for a total of more than 17,000,000 underlying evidence pieces that users can navigate to build their therapeutic hypotheses.

In the last few years, the Platform has undergone a complete rebuild, aiming to streamline data integration and harmonisation, expand users’ exploration of the data, and improve the user experience. In this context, AOTF can be seen as one of the main enhancements to the Platform user interface.

## Results

### Association on the Fly feature highlights

The Associations page is the core of the Open Targets Platform web interface, providing a comprehensive view of the evidence for a given target-disease association, ranked by overall association score.

AOTF is a revamp of this page with new, more interactive facets and additional built-in functionalities to empower users with additional control over their experience using the Platform (Fig 1a). The new feature includes a number of new applications:

**Fig 1:**
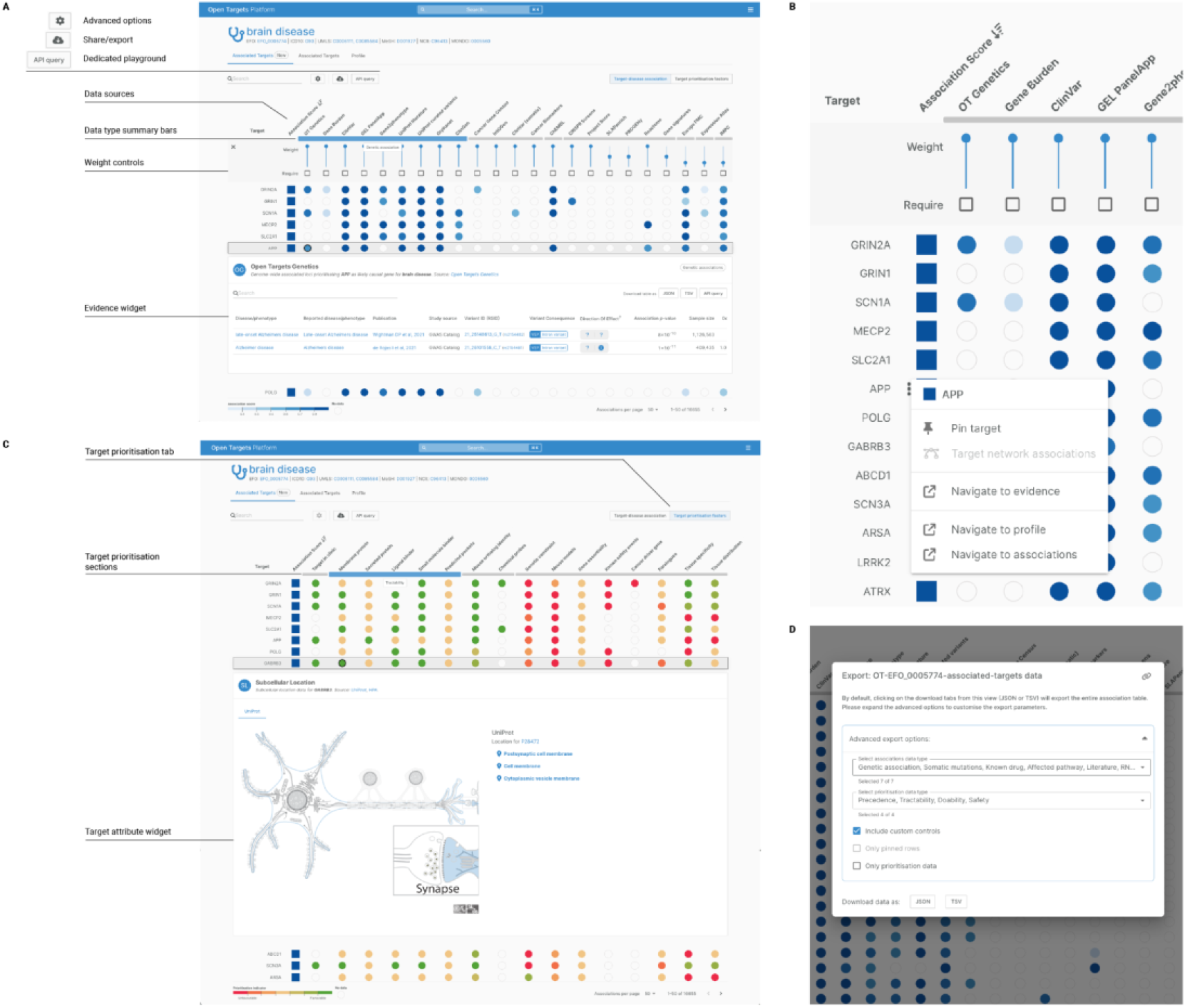
Summary panel of Associations on the Fly user interface and its subfeatures. A: Example Association on the Fly view for brain disease [https://platform.opentargets.org/disease/EFO_0005774/associations], showing all targets associated with the disease, sorted by decreasing association score (from darker to lighter blue). The main features are labelled. B: Context menu, facilitating navigation through the different Platform pages for that specific target plus target pinning (APP example). C: Example target prioritisation view “traffic light” system for target attributes [https://platform.opentargets.org/disease/EFO_0005774/associations?table=prioritisations], with darker green as most favourable moving through to deeper red as the least favourable (with arrows annotation). As highlighted in the text, a target located in the cell membrane or plasma membrane will be green in this view as it is likely to be more accessible to a drug and thus favourable (KCNB1 example). D: Export functionality, allowing users to download the entire dataset visualised in Associations on the Fly (default status) as well as customised dataset (advanced custom controls).

1. Intuitive identification of which data sources support a given disease/target association. This has been achieved by showing each data source as individual columns; data type groupings are accessible by hovering over the bar under the data source header.
2. Rapid comparison of evidence for different associations: clicking on any of the evidence buttons opens the relevant widget, enabling quick comparison of the supporting datasets, and enabling the user to rapidly sort through the evidence for a target-disease association.
3. The key feature of AOTF is the ability for users to easily modify the weighting (or relative importance) of contributing evidence from each data source. This functionality can be accessed through an advanced options tab on the user interface, additional fine-tuning is possible through an API playground. The view automatically recomputes (on the fly) the association scores based on the new user-defined weights, initiated by modification of the relevant API data sources ‘**weight’** variable on the back-end.
4. Possibility to filter by data type and data source (OR filter) by clicking on the data type summary bars or ‘required’ tick boxes.

A context menu, accessible from each entity row, has been built to allow pinning of users’ favourite targets and diseases while providing easy access to profile, association and evidence pages (fig 1b).

AOTF has been scoped, designed, and implemented with a user-centred vision. Through building user stories and user-tailored UX interviews, the Open Targets community has had a massive influence on the staged implementation and delivery of this product. For example, this feature fulfils a user-requested use case to prioritise and score target-disease associations using genetic evidence only. This is now possible by changing the weights of the non-genetics data sources to 0. Moreover, now users can also select ‘must have’ evidence from a particular data source by clicking on the corresponding box from the advanced options panel.

### Associations on the Fly subfeatures

AOTF was conceived and implemented as a reliable and dynamic feature, and this has empowered us to incorporate a second level of analysis: the Target Prioritisation factors view. (Fig 1c).

This view displays target-specific properties relevant to target prioritisation for drug discovery, individually scored, aiming to facilitate high-quality target recommendations.

Target properties are grouped into four sections: clinical precedence, tractability, doability, and safety. The scores for each target property are presented as a traffic light system: green signifies that this aspect of the target is likely beneficial for drug discovery (favourable attributes), while red indicates potential issues (unfavourable attributes). For example, a target located in the cell membrane or plasma membrane is more easily accessible to a potential therapeutic agent, so targets known to be located in the membrane will be green in this view (see example in fig 1c).

The scoring method applied to each data source is detailed in our documentation, and was developed through an Open Targets project [https://platform-docs.opentargets.org/target-prioritisation].

In order to facilitate Associations on the Fly/Target Prioritisation factor data access and analysis, we subsequently designed and developed an export functionality, allowing users to download the entire dataset (default status) as well as customised dataset — including custom controls changes, subset of data types — in tsv and json formats (Fig1c).

## Methods

Our primary goal in creating AOTF was to build a tool that would facilitate multiple levels of analysis to be represented on the same screen. For this, our initial interest was to display all the scores by data source and then explore the evidence from a specific data source. Doing it on a single screen allowed us to compare and navigate the data simultaneously without making the user switch context.

### Methodological Approach and User-Centred Design

Our web application development process followed the Agile methodology (Meso and Jain, 2006), specifically employing the Scrum framework (Cervone, 2011).

This iterative and incremental approach enables flexibility, continuous improvement, and close collaboration among team members throughout the development lifecycle. This framework has positively impacted our implementation, allowing the project team to gather feedback throughout the process, prioritise new features or adjustments, and iteratively improve the application based on user needs and market dynamics.

The process also followed a User-Centred Design (UCD) (Solano-Roman *et al*., 2019), placing a strong emphasis on understanding both the Open Targets academic and pharmaceutical communities’ needs, preferences, and behaviours throughout the design and development lifecycle. Based on research and a series of UX interviews with a pool of target users, we created detailed user personas.

These personas represented diverse user segments, as well as their goals, motivations, and user flows. We then developed journey maps, and low-fidelity wireframes to visualise and understand the typical paths users take when interacting with the web application.

This allowed us to iterate quickly and gather early input from stakeholders and users. Based on feedback and iterative design sessions, we ultimately created high-fidelity interactive prototypes using Figma [https://www.figma.com/prototyping/]. These prototypes simulate user interface interactions, helping stakeholders and users visualise the final product.

### Data processing and access

Data integration into AOTF did not involve any significant data processing mechanism, due to the intrinsic nature of our Open Targets API, which allows — through GraphQL [https://graphql.org] — access to association and evidence data. Using the methods ‘associatedTargets’ when setting a disease, or ‘associatedDiseases’ when setting a target, we obtained all the necessary data to build the Associations on the Fly. When setting a disease, we obtained extra data to generate the Target Prioritisation factor subview.

The GraphQL API was modified with the inclusion of the request of the data source ‘weight’, which was necessary to modify the order of the returned associations in the view “on the fly”, based on the user’s input (Krol, 2019).

### Tool development and integration into the Platform

We have developed this feature by using multiple open-source libraries at different application levels. The primary data component is the Open Targets API, which exposes a GraphQL API that, using Apollo GraphQL, allows us to consume the data and apply complex filtering, data source sorting, and evidence pinning.

The key libraries used in visualisation tool development are ReactJS [https://react.dev/] and TanStack Table [https://tanstack.com/table/latest]. TanStack Table allowed us to build a table while providing the flexibility to integrate customizable sections, management of states and complex transitions. ReactJS, on the other hand, facilitated presentation of the TanStack Table output through rendering and reuse of the evidence widgets already present in other parts of the Open Targets Platform. For tool deployment, we reused the same path the Open Targets Platform currently uses, by exposing an NGNX [https://nginx.org/] web server to the public using Google Cloud Platform [https://cloud.google.com/] — which was easily achieved through the Open Targets code delivery process.

The integration with the current version of the Open Targets Platform was facilitated by the use of existing tools supporting the display of evidence widgets hosted on the target-disease page. These were made accessible by a refactoring effort that, using ReactJS and Turbo [https://turbo.build/], enabled us to create a sections package and expose it to AOTF. We highlight that, to maintain good web development practices, it was necessary to create a mechanism that provides access to widgets through lazy loading. This ensures that the user is not affected by a slow loading of assets when entering the tool.

### Tool limitations

The AOTF tool has some limitations, such as the association between the data schema and the application: there is no connection between the AOTF JSON configuration and the data generated by Open Targets. Rather, the codebase hosts an object that is “hardcoded” with the desired data sources that will be displayed as columns in the user interface.

Additionally, the feature is not accessible outside of a browser or without an internet connection, and, due to a conscious decision to facilitate the development process, the tool is currently only suitable for desktop-size screens. Furthermore, the URL maximum length—which is variable within browsers— does not scale as a state management element and we are working to facilitate this in the near future.

## Conclusion and next steps

Associations on the Fly and its subview, the Target Prioritisation factor view, have been conceived with a very modular core, and this allows relatively easy planning for the next implementation steps.

Firstly, we plan to plug in an upload functionality, providing the option to upload a custom list of targets or diseases onto the two new association views. Due to the complexity and heterogeneous nature of gene and disease names, we are designing an application that will be able to suggest potential matches between the entities in the list and the ones already indexed in the Platform (Platform entity ids). When this happens, a ‘Pin hits’ tab will prompt the Associations on the Fly page to build up a custom view with the entities from the custom list.

Furthermore, following up from community requests, we plan to enable access to target interactions information within the main “Associations on the Fly” view. The interactors subview will be prompted by one of the context menu options and will contain target molecular interactions from the current Platform data feeds (String, Signor, and Reactome), sorted by their individual target-disease associations score.

These features are key to enable more complex therapeutics hypotheses building, while enhancing the Open Targets Platform user experience.

## Acknowledgements

We would like to thank the Open Targets Core Team, including past members, for their contribution to the Open Targets Platform. We would also like to thank our partners, collaborators and users who have helped with the UX sessions.

## Bibliography

1. Ochoa, D. et al. The next-generation Open Targets Platform: reimagined, redesigned, rebuilt. Nucleic Acids Res. 51, D1353–D1359 (2023).

2. Meso, P. & Jain, R. Agile Software Development: Adaptive Systems Principles and Best Practices. Inf. Syst. Manag. 23, 19–30 (2006).

3. Cervone, H. F. Understanding agile project management methods using Scrum. OCLC Syst. Serv. Int. Digit. Libr. Perspect. 27, 18–22 (2011).

4. Solano-Roman, A. et al. ‘NX4: a web-based visualization of large multiple sequence alignments’, Bioinformatics. Edited by A. Valencia, 35(22), pp. 4800–4802 (2019).

5. Krol, K. ‘Handling Data Sources and Prioritization in GraphQL APIs’, Journal of Web Engineering, 18(2), pp. 167–180 (2019).

